# MeXpose - A modular imaging pipeline for the quantitative assessment of cellular metal bioaccumulation

**DOI:** 10.1101/2023.12.15.571675

**Authors:** Gabriel Braun, Martin Schaier, Paulina Werner, Sarah Theiner, Jürgen Zanghellini, Lukas Wisgrill, Nanna Fyhrquist, Gunda Koellensperger

**Author notes:** These authors contributed equally to the main findings of this manuscript (Gabriel Braun, Martin Schaier). Corresponding author: Gunda Koellensperger Institute of Analytical Chemistry, 1090 Vienna, Austria.

## Abstract

We introduce MeXpose, an imaging pipeline for single-cell metallomics by laser ablation inductively coupled plasma time-of-flight mass spectrometry (LA-ICP-TOFMS). MeXpose is designed for mechanistic studies on metal exposure unravelling cellular phenotypes and tissue level characteristics of metal bioaccumulation. MeXpose leverages the high-resolution capabilities of low-dispersion laser ablation setups, a standardised approach to quantitative bioimaging, and the toolbox of immunohistochemistry using metal-labelled antibodies for cellular phenotyping. MeXpose further offers the full scope of single-cell metallomics via an extended mass range accessible through ICP-TOFMS instrumentation (covering isotopes from m/z 14-256) and integration of a complete image analysis workflow. This enables studying quantitative metal accumulation in phenotypically characterized tissue at cellular resolution. Metal amounts in the sub-fg range per cell can be absolutely quantified. As a showcase, an *ex vivo* human skin model exposed to cobalt chloride (CoCl_2_) was investigated. Metal permeation was studied for the first time at single-cell resolution, showing high bioaccumulation in the epidermal layers and especially in mitotic cells, accumulating cobalt (Co) in the low fg range per cell. In this cellular phenotype, Co accumulation was correlated to DNA damage. While the amount of cobalt was significantly lower in the collagenous matrix of the dermal layer, cells in the vicinity of blood vessels and smooth muscle showed significant Co deposits as well. MeXpose provides unprecedented insights into metal bioaccumulation with the ability to explore novel relationships between metal exposure and cellular responses on a single-cell level, paving the way for advanced toxicological and therapeutic studies.

**Graphical abstract:** 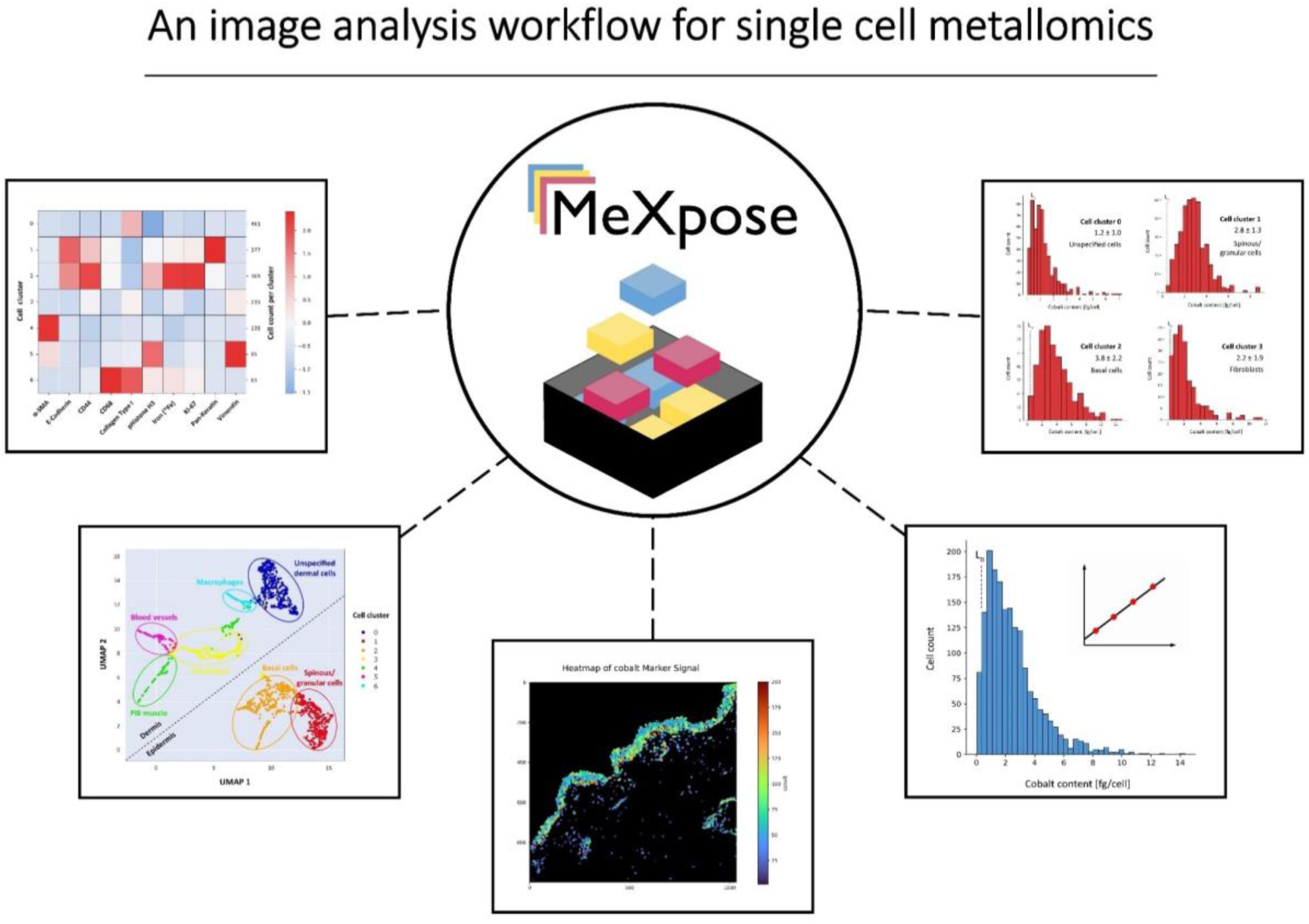

## Introduction

Metal exposure in human populations is increasing, driven by anthropogenic activities, modern industrialization and medical use. For a significant proportion of metals (and metal-containing molecules), exposure is accompanied by bioaccumulation in various tissues, with implications on human health^1, 2, 3, 4^. The toxic effect of metals depends on the chemical form, dose and route of exposure. Accumulation patterns of metal species differ significantly among various tissue- and cell types^5, 6^. More importantly, metal exposure influences dynamic cellular properties, thereby potentially modifying the behaviour of cellular metal accumulation^7, 8^. For example, cancer cells exposed to cytotoxic/cytostatic metal compounds may acquire resistance by developing a distinct cellular phenotype characterized by altered metal accumulation behaviour^9, 10, 11, 12^. In keratinocytes, cobalt accumulates in the cell nucleus and in perinuclear structures, causing multiform toxicity related to interaction with genomic DNA and nuclear proteins, as well as to altered zinc and magnesium homeostasis.^13^

Therefore, we hypothesize that metal activity and patterns of toxicity could be better understood by measuring the characteristics of metal tissue distribution at single-cell resolution. Retaining spatial information is essential, as metal concentration gradients within the tissue arise upon routes of exposure and cell activities are affected by position dependent interactions and nutrient exchange.

Assessing metal accumulation in single cells is a challenging analytical task^14, 15, 16, 17, 18, 19^. Time-resolved inductively coupled plasma mass spectrometry (ICP-MS) measurements enable the quantification of metals in thousands of cells at high throughput. The underlying analysis principle involves the production of single-cell events in a flow cytometer from cell suspensions which are subsequently detected by ICP-MS^20, 21^. As a drawback, the tissue context is not retained. More importantly, proof-of-principle studies focusing on metal quantification miss multiplexing measurement capability by limiting the single analysis to one element per cell^12, 22, 23, 24, 25, 26, 27^ . Thus, metal uptake quantified at the single-cell level can only be correlated to predominant cellular bulk properties, such as the cell cycle in synchronized cell cultures.^28^ Multiplexed analysis of single cells was achieved upon introduction of time-of-flight-based ICP-MS instruments, equipped with a quasi-simultaneous mass analyser recording an entire spectrum of elements from a single ion pulse^20^. When combined with antibody labelling strategies using metal isotope tags, the mass spectrometric pendant of fluorescence-based cytometry boosted the number of read-outs per cell, allowing in-depth phenotypic screenings of cell populations^21, 29^. Up to date, rare examples of mass cytometry studies link cellular properties to metal accumulation at the single-cell level^30, 31, 32^.

The imaging version of mass cytometry combines laser ablation (LA) using low-dispersion setups with ICP-TOFMS allowing single-cell analysis with spatial resolution down to 1 µm^33, 34,35^. Imaging mass cytometry (IMC) creates unprecedented molecular and cellular maps^36, 37, 38^.

Characterization of cellular features is supported by tailored image analysis pipelines ^39, 40, 41, 42, 43^. Many of these solutions are subject to continuous improvements, since existing pipelines show a combination of various shortcomings, such as a) inability to provide a complete end-to-end data analysis workflow; b) high level of required technical entry knowledge; c) lack of interactivity or scalability. Most important, none of the current available image analysis solutions contain the functionality to provide quantified absolute metal contents for tissue imaging.

While recent standardization strategies^44, 45, 46, 47, 48^ improved quantitative bioimaging by LA-ICP-MS, absolute quantitative values are primarily assessed on a pixel basis, or for regions of interest^49, 50, 51, 52^, Cell-based metal abundances (fg per cell) can only be derived upon integrating cell segmentation in the image analysis pipeline, which in turn requires adequate spatial resolution (down to 1 µm for single cell analysis in tissue) and multiplexed measurements. As a minimum prerequisite, at least one membrane and one nucleus marker as well as the accumulating metal must be measured simultaneously. The majority of quantitative bio-imaging studies fails to fulfil all technological requirements, impeding quantitative single cell analysis in tissue.

We introduce MeXpose, an LA-ICP-TOFMS imaging pipeline, linking quantitative metal distribution at single-cell resolution with identification of cellular phenotypes in tissues. MeXpose exploits the full scope of IMC, identifies cells preferentially accumulating metals, and quantifies their metal content. While IMC was established on an ICP-TOFMS instrument with a truncated mass range (m/z>75 Da)^20, 21^, necessary to avoid saturation effects on the detector, the ICP-TOFMS instrumentation fundamental to MeXpose offers an expanded mass range towards lower masses (m/z range 14-256). It successfully removes argon ions, otherwise negatively impacting on the sensitivity, by placing notch filters in the ion optics^53, 54, 55^. Thus, potential toxic metal species such as Co, Cr, Ni, Ti species can be investigated. Another key component of MeXpose is the quantitative bioimaging capability. Each imaging experiment is accompanied by a day-to-day fully traceable calibration routine^46^. MeXpose integrates for the first time this quantification strategy and IMC into one stringent end-to-end workflow. By utilizing metal-labelled antibodies, MeXpose can determine the metal content of segmented cellular objects in tissue samples. Beyond merely locating the metal-accumulating cells in tissue, these cells are phenotypically characterized. This way, MeXpose paves the way for novel research on metal toxicity, biology and medicine, aiming to provide a mechanistic understanding of metal activity for both “wanted” and “unwanted” metal exposures.

The MeXpose toolbox is showcased in a proof-of-concept study, showing the power in mechanistic exposure studies on metals in *ex vivo* human skin tissue. Several metals such as Ni, Cr, Co and Au are known to elicit contact allergies upon direct skin contact in sensitized individuals^56, 57^. There is no clear-cut correlation between sensitizing potency of different metals, their binding kinetics and the resulting penetration depth into the skin layers, indicating that the mechanisms of skin penetration, as a first step in contact allergy, might be more complex than assumed^58, 59^. In this work, CoCl_2_, a clinically highly relevant metal cation with proven skin penetration capacity was selected and studied by MeXpose in an *ex vivo* human skin model^58, 60, 61^. For the first time, studies on cobalt permeation in human skin include the quantification of metal uptake of single cells in tissue, together with their spatial distribution and phenotypical characterization. In fact, MeXpose uniquely discovered distinct cellular phenotypes and tissue locations preferentially accumulating cobalt once the metal penetrated the skin barrier.

## Results

### Step by step- from measurement to data analysis

MeXpose relies on low-dispersion LA offering single-cell resolution (1 µm pixel size) and pixel acquisition rates up to 1 kHz in combination with multi-element ICP-TOFMS analysis^62, 63, 64^. The ICP-TOFMS system provides a mass coverage of m/z = 14-256, enabling the study of metal exposure of elements from the lower mass range such as Cr, Co and Ni and the cellular metallome including Fe, Cu and Zn^65^. Tissue sections are stained with metal-conjugated antibodies and measured in a sequence with multi-elemental calibration standards based on gelatine micro-droplets, dispensed on surfaces by robotics^46^. The droplets feature precisely characterized pL volume and (multi-)elemental concentrations. Provided complete ablation of the micro-droplet is achieved, an accurate elemental amount per micro-droplet can be inferred and used for constructing an external calibration function. The calibration sequences, consisting typically of a blank and 5 standards of different concentration levels can be measured within few minutes by LA-ICP-TOFMS. The linear dynamic working range typically covers 2 orders of magnitude (fg amounts). As in any quantification exercise using ICP-MS, the standardization measurement is performed on a day-to-day routine. The acquired data is then evaluated using MeXpose, which provides comprehensive tools for single-cell analysis. Figure 1 gives an overview of the entire analysis procedure.

**Figure 1:**
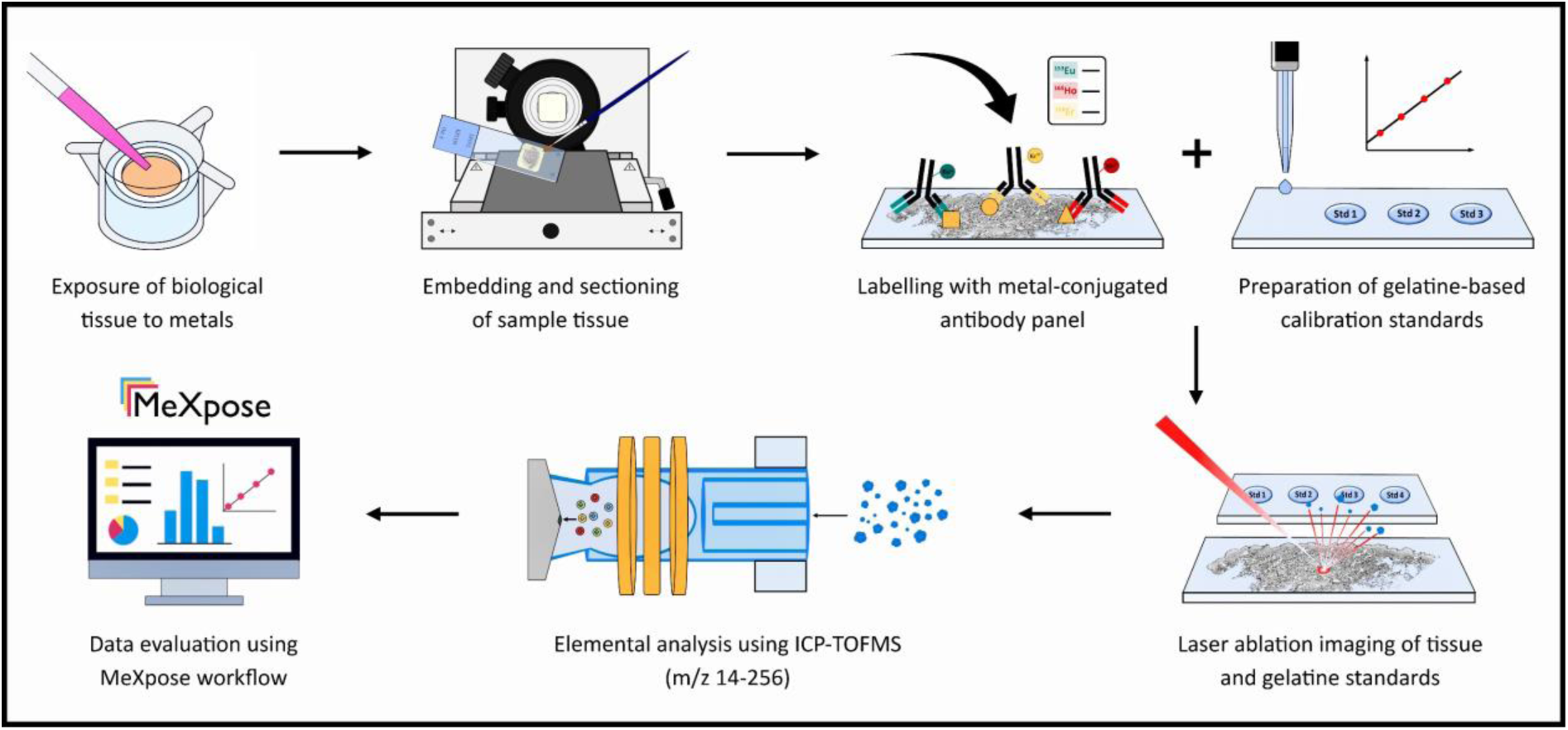
Overview of the workflow to assess metal exposure at the single-cell level. Metal accumulation experiments are carried out on biological tissues. The tissues are embedded in FFPE (formalin-fixed, paraffin-embedded), sectioned, and labelled with a panel of metal-conjugated antibodies. The samples, and subsequently the multi-element calibration standards, are subjected to LA-ICP-TOFMS analysis. Data analysis is performed using the MeXpose workflow.

### MeXpose data analysis

MeXpose deploys a dedicated high-dimensional multiplexed image analysis pipeline combining various third-party software tools with additional newly developed Python scripts into a user-friendly, flexible containerized platform. Figure 2 outlines the workflow and key features. The elemental images obtained by LA-ICP-TOFMS are exported as single- or multi-channel tiff format and submitted to the technology agnostic workflow. MeXpose data evaluation offers both, an interactive and a scalable semi-automated version deployed as a single docker container. The latter was developed to meet the requirements of large-scale studies. The interactive mode was designed to enable immediate control on pre-processing, segmentation and exploratory data analysis. This approach ensures optimal cell segmentation for different tissue types, which is a key prerequisite for quantitative image analysis.

**Figure 2:**
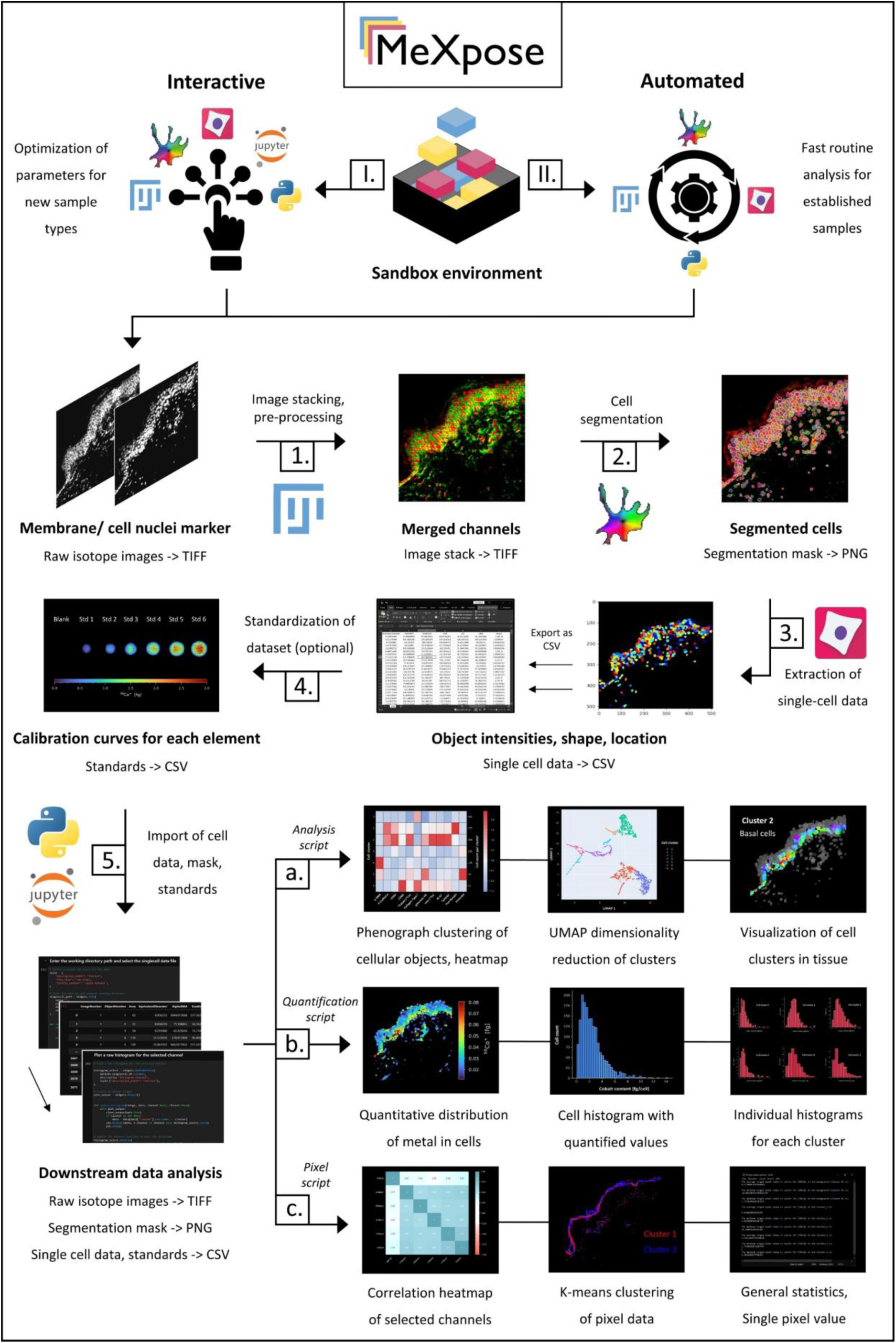
MeXpose - a pipeline for image analysis at the single-cell level. User defined elemental channels are stacked, pre-processed, and used to run a pre-trained network to generate cell masks. Consequently, a feature matrix is extracted from the raw data using the segmentation mask. The extracted data can be used for image analysis at the single-cell level by downstream statistical analysis. The quantitative module of MeXpose integrates an external calibration function, which can be applied on the data to obtain metal quantities per cell.

Depending on the individual needs, all included applications can be executed in ‘GUI-mode’ (Graphical User Interface) by simply launching them using ‘-gui’ as a suffix to the desired applications name. The core functionality of both workflows (interactive and scalable) is the same and utilises identical software with the exception being that the interactive Jupyter notebooks are replaced by command line python scripts in the automated workflow.

Both MeXpose versions – i.e. the interactive and scalable workflow - offer essential functionalities to obtain tabular cellular data characterized by object-based integrated channel/isotope intensities and morphology for downstream quantitative analysis. For data pre-processing, cell segmentation, exploratory data analysis and visualization, state-of-the-art software packages dedicated to each individual step were combined in a modular end-to-end pipeline. MeXpose deploys the Fiji (v1.54f) image processing software for pre-processing and multichannel image visualisation, Cellpose 2.0 (v2.2.3) for deep learning assisted cell segmentation, Cellprofiler (v4.2.1) for feature extraction (intensities and morphology) of cellular objects and newly developed in-house Python scripts (v3.8.8) and JupyterLab notebooks (JupyterLab v3.5.3; Jupyter Notebook v6.5.6) for data analysis and visualisation^66, 67, 68, 69, 70^. Each module can be replaced with other software at will, which allows for seamless integration into existing image analysis workflows. The obtained tabular single-cell data are exported for further quantitative image analysis. MeXpose integrates scripts for absolute metal quantification and for statistical analysis. The latter task involves established strategies of image analysis, such as unsupervised clustering and/or dimensionality reduction. Phenograph (v1.5.7), an unsupervised graph clustering algorithm based on the Leiden algorithm, is used to categorise segmented cells into clusters depending on their feature intensities and inter-versus intra-cluster connections^71^. Simplified, when represented as a graph, cells that form a subpopulation with a high number of internal connections and a lower number of external connections to other subpopulations are designated as a cluster. MeXpose allows to visualize these high dimensional clusters by projecting them as a two-dimensional embedding using UMAP (v0.5.4)^72^. This enables visual inspection of cluster quality. By creating a cluster heat map, MeXpose permits in-depth analysis of the channel expression profile and relative intensities within each cluster, enabling characterization of the various cellular phenotypes within tissue. Data interpretation is facilitated by the MeXpose option of projecting the corresponding segmented objects onto elemental images as a channel heatmap for visual inspection of spatial arrangements.

MeXpose uniquely allows to apply a calibration function to the obtained data (including pixel-based and cell-based data) resulting in object-based quantitative integrated elemental amounts. Following standardization, the colour intensity code in the images can be converted to a colour concentration code. Upon visualizing segmented cells on the images with a colour concentration code, quantitative bioaccumulation resulting from metal exposure can be observed at the single-cell level. The images showing quantitative single-cell metal data can be generated for a user-defined cell population. Finally, the single-cell metal quantities can be plotted as histograms either for the complete imaged cell populations, or again for certain sub-populations, e.g. cell phenotypes as revealed by statistical analysis. Producing histograms and plotting quantified cellular objects for distinct cellular sub-populations greatly aid the discovery, of so far missing links between cellular metal enrichment and cellular phenotypes and/or tissue architecture.

### Revealing the structural landscape of human skin through multiplexed imaging

To make the case for single-cell metallomics, CoCl_2_ permeation in human skin was studied for the first time at single-cell resolution. Skin represents one of the major metal exposure routes in occupational settings. A cutting-edge *ex vivo* human skin model, NativeSkin^®^ was exposed to CoCl_2_. The model closely resembles *in vivo* skin permeation upon exposure, as it preserves tissue architecture in controlled environments^73^. Figure 3 shows LA-ICP-TOFMS images of the *ex vivo* skin model obtained by overlay visualizations of selected image channel intensities. The overlay images unveiling the skin tissue structural architecture relied on five markers along with one marker for cell nuclei (6 channels). The selected antibody panel (Supplementary Table 1) characterized the different layers of the epidermis and the dermis together with essential structural features such as blood vessels, sweat glands and muscle tissue.

**Figure 3:**
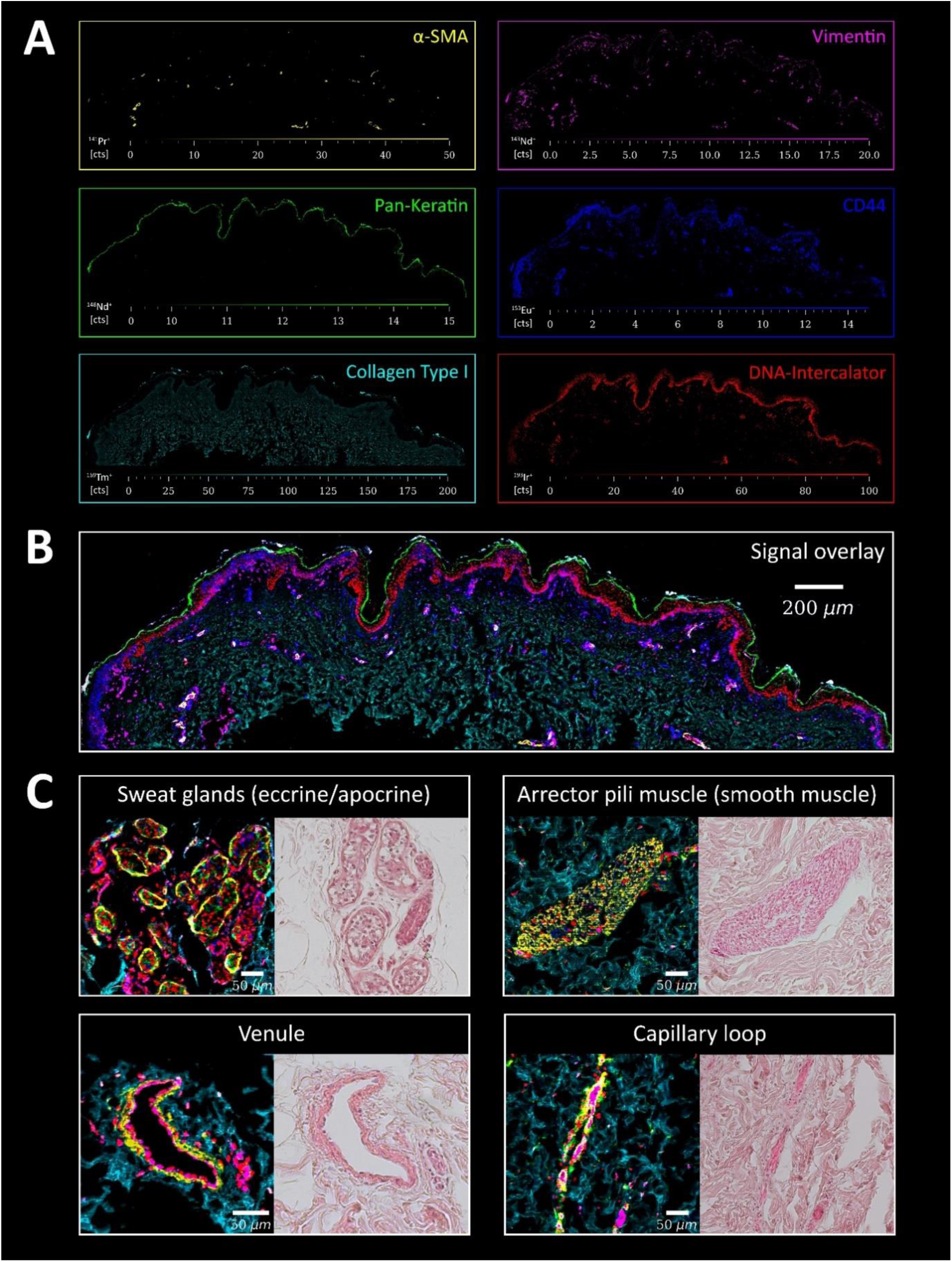
Skin tissue architecture. Following FFPE treatment and sectioning, the skin thin sections were labelled with metal-conjugated antibodies and analysed by LA-ICP-TOFMS (1 µm pixel size, 200 Hz pixel acquisition rate). The applied histochemistry and the DNA intercalator allowed to visualize the epidermal and dermal layers together with essential structural features, including blood vessels, sweat glands, and muscle tissue. The marker panel included α-SMA (specifically labels smooth muscle cells), vimentin (mesenchymal cells, encompassing fibroblasts of the papillary dermis, as well as the endothelium of blood vessels), pan-keratin (cornified layer and keratinocytes in the epidermis), CD44 (cell surface receptor for hyaluronic acid, membrane marker for both epidermal and dermal cells), collagen (extracellular matrix of the dermis) and DNA intercalator (cell nuclei). (A) shows the signal intensity maps of the individual markers, (B) visualizes an overlay of the 6 channels, (C) shows structural features of regions of interest of the skin tissue together with H&E staining of consecutive FFPE sections for comparison.

### Single-cell analysis in human skin tissue

Following the interactive branch of the MeXpose image analysis pipeline, single-cell data were extracted for human skin tissue sections. Stacking required the selection of two channels, (1) a selective membrane and (2) a nuclei marker. In human skin, CD44, proved to be a valuable membrane marker, while the established Ir-based DNA intercalator was selected as nuclei marker (Supplementary Fig. 1). Before stacking, the two channel images were pre-processed using a combination of steps, including contrast enhancement and intensity thresholding, outlier filtering and median/gaussian filtering. Mexpose deploys Cellpose 2.0 for cellular segmentation, leveraging its integrated pre-trained ‘model zoo’ as well as its interactive GUI (interactive workflow only). It also inherently includes segmentation quality control through visualisation and the ability to interactively correct segmentation errors and add custom annotations. Upon completion, Cellpose generates PNG cell masks which are successively used to extract multi-parametric tabular single-cell data (characterized by morphology and integrated intensities) from raw images using Cellprofiler.

MeXpose image analysis of human skin areas in the mm^2^ range, revealed cell numbers in the 10^3^ orders of magnitude with an average cell diameter of 10 µm. Supplementary figure 1 shows the obtained segmented cells in the human skin model, projected on top of the LA-ICP-TOFMS image. High cell numbers were observed in the epidermis, while the dermal layers showed fewer cells embedded in the collagenous matrix.

The applied downstream statistical analysis resorted to a marker panel of 9 metal-conjugated antibodies (Supplementary Table 1) and the endogenous metal iron. Phenograph clustering revealed seven clusters, which could be assigned to cellular phenotypes, typical for the different epidermal and dermal layers. Keratinocytes (basal, spinous and granular), macrophages, fibroblasts and smooth muscle cells could be assigned (Fig. 4). MeXpose uniquely allows the consideration of endogenous elements upon cellular phenotyping. In the case of human skin, adding cellular iron to the marker panel was beneficial for characterizing epidermal keratinocytes. Basal cells, which have the highest blood flow due to their proximity to capillaries, could be differentiated from other keratinocytes using iron (Supplementary Fig. 2)^74, 75^. Larger blood vessels such as arteries and veins, sweat glands and hair follicles also showed characteristic iron distributions that are useful for cell pheno-/subtyping (Supplementary Fig. 3).

**Figure 4:**
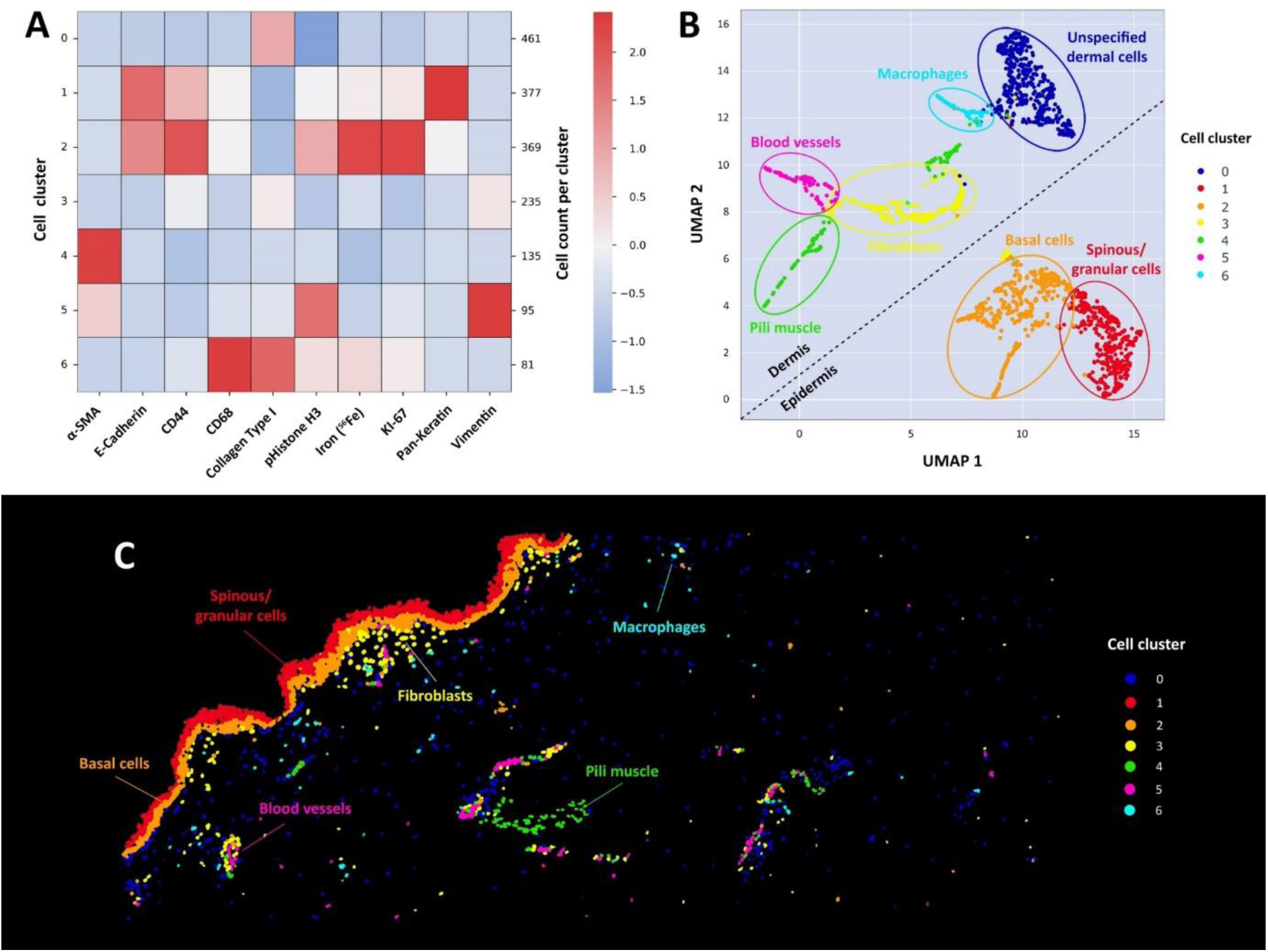
Downstream statistical analysis. Using the marker panel collagen type-I, Pan Keratin, E-Cadherin, CD44, Vimentin, α-SMA, CD68, KI-67, pHistone H3 and endogenous iron revealed 7 distinct cell populations (0-6). (A) Heatmap and (B) UMAP are shown for the 1782 segmented cellular objects. Cluster 1 and Cluster 2 relating to the epidermis were well separated from the dermal clusters (0, 3-6). (C) Visualisation of the Phenograph clusters in the segmentation mask, revealing the location of the cells within the skin tissue.

### Quantifying metal accumulation at the single-cell level

The human skin model exposed to CoCl_2_ showed pronounced metal permeation into the epidermis. High bioaccumulation was found down to a depth of approximately 100 µm, however, the highest Co amount was observed within the cornified layer at a permeation depth of 20 µm (Supplementary Fig. 4). Only small amounts of cobalt were found to reach the dermal layer of the skin. By applying the MeXpose external calibration function (obtained from the measurement of gelatine based micro-droplets) (Fig. 5A), both pixel-based and cell-based intensity data on cobalt were converted into absolute amounts (Fig. 5B, C). Within the MeXpose pipeline, the obtained single-cell quantities can be plotted as histograms showing the distribution of cellular metal content for subpopulations or the entire cell population. For visualization, projections of segmented cells on top of the image are associated with a colour coded concentration scale. The entire imaged cell population of 1782 cells revealed a mean Co concentration of 2.4 fg per cell (Fig. 5D). A procedural limit of detection of 0.38 fg Co per cell was estimated, by studying control skin tissue samples (not exposed to CoCl_2_) that were stained with the metal-conjugated antibody panel (Supplementary Fig. 5). The cellular Co intensities constituting the blank values were converted into quantities and treated as Poisson distributed to infer the limit of detection^76^. To prove the method’s fit for purpose, additional validation experiments were carried out. Two consecutive skin tissue sections were exposed to Co and measured (1) following immunostaining and (2) omitting the sample preparation steps (Supplementary Fig. 6). Comparative pixel-based image analysis showed the validity of the quantitative bioaccumulation data.

**Figure 5:**
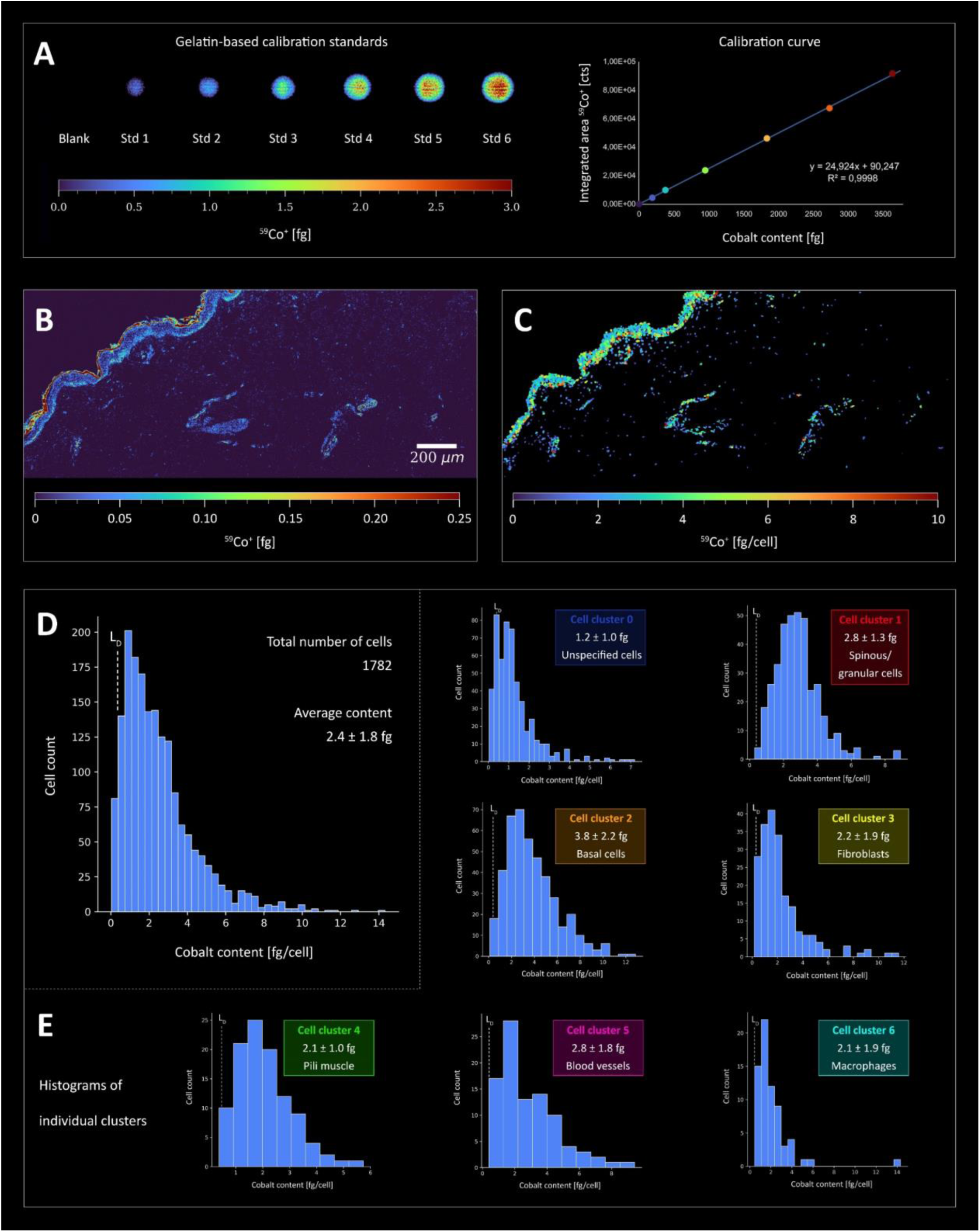
Quantitative Co imaging in human skin at the single-cell level (LA-ICP-TOFMS analysis, 1 µm pixel size, 200 Hz pixel acquisition rate). *Ex vivo* skin models were incubated with 100 µg cm^-2^ CoCl2 solution (dose chosen based on previous results from patch tests in Co allergic subjects) for 48 h. (A) Quantification of Co was performed using gelatine-based calibration standards which were measured along with the skin thin sections. (B) The Co signal intensity map of the tissue was converted to absolute Co amounts using the calibration factors. Co was enriched in the cornified layer, but penetrated the epidermis, whereas the dermis showed significantly lower Co quantities. (C) Pixel data were transformed into cellular data using MeXpose, to better visualise the Co accumulation within cells. The histogram in (D) shows the cobalt distribution in all segmented cells. The average Co amount per cell is 2.4 ± 1.8 fg. Panel (E) shows histograms for each individual cell cluster revealed by unsupervised clustering (Fig. 4). Cluster 2 (basal cells) in particular showed significantly higher Co content compared to the other cell types (Supplementary Fig. 7).

### Cellular phenotypes of cobalt accumulation

The human skin characteristics of Co bioaccumulation were investigated by assessing the quantitative single-cell Co data and their distribution for the observed cell phenotypes, respectively (Fig. 5E). Among the 7 sub-populations obtained from unsupervised Phenograph clustering, mitotic basal cells of the epidermis (cluster 2 in Fig. 4) showed pronounced Co bioaccumulation, with an average of 3.8 fg/cell. When applying the null hypothesis (Mann-Whitney U), this sub-population was shown to differ significantly from other clusters (Supplementary Fig. 7). Besides basal cells, spinous and granular cells (cluster 1) showed high metal bioaccumulation. In these epidermal cells, high Co levels correlated with an increase in DNA damage (Supplementary Fig. 8). Co accumulation was also pronounced in vascular endothelial cells (Supplementary Fig. 9), with an average of 2.8 fg/cell, particularly in capillaries of the papillary dermis characterised by loose connective tissue (Supplementary Fig. 10). Clusters associated with arrector pili smooth muscle cells and fibroblasts showed comparatively lower Co accumulation (clusters 3 and 4). Macrophages in cluster 6 showed a similar distribution with an average cobalt content of 2.1 fg/cell. The remaining “unspecified” cell population of the dermis, not characterised by the selected antibody panel (cluster 0), showed the lowest cobalt bioaccumulation (mean value of 1.2 fg/cell).

## Discussion

MeXpose introduces cutting edge single-cell metallomics in tissue, combining quantitative bioimaging by LA-ICP-TOFMS and the comprehensive toolbox of imaging mass cytometry into an integrated end-to-end workflow. MeXpose couples comprehensive tissue characterisation and cell population assessment with quantitative cellular metal bioaccumulation analysis. Additionally, the MeXpose standardization concept allows to infer absolute amounts of metals on a single-cell level in tissue. As any quantification exercise, MeXpose quantitative assessment of bioaccumulation calls for validation, which has to be performed for each metal species and tissue type, respectively. Generally, the method is fit-for-purpose for metals forming macromolecular complexes in tissue upon exposure. Metal accumulation in the sub-fg range per cell can be studied.

The prime example of Co exposure and its effects, studied in NativeSkin^®^ models, demonstrated the value of the approach. Co skin permeation was investigated at unprecedented detail, i.e. at the single-cell level. The application of cobalt in the form of CoCl_2_ to the skin resulted in a predominant accumulation in the cornified layer, a key skin barrier. However, our investigation also revealed a notable ability of Co to penetrate deeper layers of the skin, primarily targeting the epidermis and, to a lesser extent, the dermis. This transdermal permeation resulted in significant Co accumulation within the basal layer, correlating with increased DNA damage at the single-cell level. Due to the close proximity of basal cells to dermal blood vessels, there is a risk that Co could be transported further into the blood stream. This concern was supported by the elevated levels of Co in vascular endothelial cells. High levels of Co in the systemic circulation could potentially contribute to organ damage due to its ability to generate reactive oxygen species.

## Methods

### Chemicals and Reagents

Ultrapure water (18.2 MΩ cm) from the ELGA water purification system (Purelab Ultra MK 2, United Kingdom) was used for all dilutions and washing steps. A multi-element stock solution was obtained from Labkings (Hilversum, The Netherlands). Bovine serum albumin (lyophilised powder, BioReagent), Tris-buffered saline (BioUltra), m-xylene (anhydrous, ≥99%) and ethanol (absolute, EMSURE®) were purchased from SigmaAldrich (Steinheim, Germany). Tween-20 detergent solution (Surfact-Amps™, 10%) and SuperBlock^TM^ blocking buffer (TBS) were obtained from Thermo Fischer Scientific (Waltham, MA, USA). Target retrieval solution (pH 9, Tris/EDTA) was supplied by Agilent Technologies (Waldbronn, Germany). Metal-conjugated antibodies listed in Supplementary Table 1 and the Ir-intercalator (Cell-IDTM, 125 µM) were purchased from Standard BioTools (San Francisco, CA, USA) and CD16/32 antibody from BD Biosciences (San Jose, CA, USA). The *ex vivo* skin model (NativeSkin^®^) was bought from Genoskin (Toulouse, France). CoCl_2_ hexahydrate (CoCl_2_ · 6H_2_O) was purchased from Merck KGaA (Darmstadt, Germany). Biopsies were taken from the residual skin of adults who had undergone surgery. Informed consent was obtained from the donors and the skin model was approved by the Ethics Committee. A detailed view of the skin model is shown in Supplementary Fig. 11.

### Exposure of *ex vivo* skin model to CoCl_2_

Upon arrival of the skin model, the media (supplied by Genoskin) was added under sterile conditions and incubated for 1 h at 37°C, 5% CO2, 95% relative humidity (according to user manual). A 100 mg ml^-1^ stock solution of CoCl_2_ · 6H_2_O was prepared by dissolving 1 g in 10 ml ultrapure water followed by sterile filtration. For the target dose of 2.5 µg µl^-1^ (based on prior results from patch testing of cobalt-allergic individuals), 25 µl of the stock solution was mixed with 975 µl of sterile ultrapure water and vortexed prior to exposure^60, 77^. The CoCl_2_ solution was added to the model by pipetting 20 µl. The solution was homogeneously distributed on the apical side of the model. The plate was then placed in an incubator (37°C, 5% CO2, 95% relative humidity). The next day the media was changed. Samples were collected after 48 h, later embedded in FFPE and sectioned at 5µm.

### Immunostaining of skin tissue sections

Skin tissue sections were deparaffinized by treatment with fresh xylene for 20 min. Descending grades of ethanol (100-70%) were used for rehydration. After rinsing with ultrapure water, heat-mediated antigen retrieval was performed at 96°C for 30 min using a Tris-EDTA buffer at pH 9. Slides were then cooled and washed with ultrapure water and TBS/0.05% Tween. To prevent non-specific binding, SuperBlock buffer was applied to the skin tissue for 30 min at RT, followed by CD16/32 treatment for 10 min. The sections were then exposed to a cocktail of metal-conjugated antibodies overnight in a hydration chamber at 4°C. Details of the metal-conjugated antibodies used are given in Supplementary Table 1. The antibodies (1:100 dilution) were prepared in a mixture of 0.5% BSA, 1:100 CD16/32 in TBS/0.05% Tween. To prevent aggregation, the antibodies were centrifuged at 13,000 g for 2 min prior to use. Tissue sections were then stained with the Ir-based DNA-Intercalator (125µM) using a 1:100 dilution in TBS/0.05% Tween. After incubation for 5 min at RT in a hydration chamber, slides were thoroughly washed with ultrapure water and air dried. Microscopic images were taken prior to LA-ICP-TOFMS analysis to provide an overview of the tissue sections.

### Calibration standards for LA-ICP-TOFMS analysis

Quantitative analysis was carried out by LA-ICP-TOFMS with the use of gelatin micro-droplets, as described previously^46, 78^. For this purpose, multi-element standard solutions were prepared gravimetrically using commercial standard stock solutions and mixed with a gelatin solution. The resulting solutions were then transferred to the wells of a 384-well plate. The plate served as the sample source for a micro-spotter system.

The CellenONE X1 micro-spotter (Cellenion, Lyon, France) was used for generating arrays of gelatin micro-droplet standards on glass slides. These droplets were approximately 200 µm in diameter and had a volume of 400 ± 10 pL. The software of the instrument evaluated the size of the droplets, which was then used for normalization to determine the absolute amounts of elements in the droplets. The entire micro-droplets were subjected to quantitative and selective ablation followed by multi-element analysis using LA-ICP-TOFMS.

### LA-ICP-TOFMS analysis

An Iridia 193 nm laser ablation (LA) system from Teledyne Photon Machines (Bozeman, MT, USA) was coupled to an icpTOF 2R ICP-TOFMS instrument from TOFWERK AG (Thun, Switzerland). The LA system consisted of an ultra-fast, low-dispersion cell in a Cobalt ablation chamber^62, 63^. The cell was connected to the ICP-TOFMS using an aerosol rapid introduction system (ARIS). An argon make-up gas flow (∼0.90 L min^-1^) was introduced through the low-dispersion mixing bulb of the ARIS into the He carrier gas flow (0.60 L min^-1^) before entering the plasma. The NIST SRM612 glass certified reference material (National Institute for Standards and Technology, Gaithersburg, MD, USA) was used for daily tuning. Tuning was aimed at high intensities of certain ions (^59^Co^+^, ^115^In^+^ and ^238^U^+^), minimal oxide formation (^238^U^16^O^+^/^238^U^+^ ratio < 2%) and low elemental fractionation (^238^U^+^/^232^Th^+^ ratio ∼ 1). In addition, aerosol dispersion was minimized by optimization of the pulse response time for ^238^U+ with the FW0.01M criterion. Laser ablation sampling was performed in fixed dosage mode 2 with a 200 Hz repetition rate. A circular spot size of 2 µm was used with a line spacing of 1 µm, resulting in a pixel size of 1 µm x 1 µm. By using energy densities above the ablation threshold of the samples but below that of glass, selective ablation of the samples was achieved^79^. Gelatin micro-droplets and skin tissue were quantitatively removed using fluences of 0.4 J cm^-2^ and 1.0 J cm^-2^, respectively. The icpTOF 2R ICP-TOFMS instrument offers a specified mass resolution of 6000 (R = m/Δm) and detects ions in the m/z range of 14 to 256. Its integration and readout rates were in line with the LA repetition rate. The instrument was equipped with a torch injector of 2.5 mm inner diameter and nickel sample and skimmer cones (with a 2.8 mm skimmer cone insert). It was operated with a radiofrequency power of 1440 W, an auxiliary Ar gas flow rate of ∼ 0.80 L min^-1^ and a plasma Ar gas flow rate of 14 L min^-1^. All measurements were performed in collision cell technology (CCT) mode, with the collision cell pressurized with a H2/He gas mixture (93% He (v/v), 7% H2 (v/v)) at a flow rate of 4.2 mL min^-1^. Detailed instrumental parameters for LA-ICP-TOFMS measurements are given in Supplementary Table 2.

### Data acquisition and processing of LA-ICP-TOFMS data

LA-ICP-TOFMS data were acquired using TofPilot 2.10.3.0 from TOFWERK AG (Thun, Switzerland) and stored in the open-source hierarchical data format (HDF5, www.hdfgroup.org). Subsequent data processing was done using Tofware v3.2.2.1 (TOFWERK AG, Thun, Switzerland), used as an add-on to IgorPro (Wavemetric Inc., Oregon, USA). This processing included three steps: (1) correction of mass peak position drift across the spectra with time-dependent mass calibration, (2) determination of peak shapes (3) fitting and subtraction of the mass spectral baseline. The data were further processed using HDIP version 1.8.4.120 from Teledyne Photon Machines (Bozeman, MT, USA). A built-in script was used to automate the processing of files generated by Tofware, producing 2D elemental distribution maps. Thus, acquired LA-ICP-TOFMS data were processed in HDIP and exported as separate TIFF files for each isotope to enable single-cell based analysis, or in CSV format for pixel-based image analysis.

### MeXpose image analysis

TIFF files or CSV files were imported for single-cell- or pixel-based image analysis, respectively.

### MeXpose implementation

MeXpose contains all the software and scripts needed to run the workflow and was designed to be used as an all-in-one Docker container. The MeXpose image can be found on the Docker Hub under the name MeXpose. It contains both the interactive and scalable versions of the workflow.

### MeXpose description

In principle, the workflow of the MeXpose data analysis consists of four steps: Data pre-processing, cell segmentation, data extraction and downstream statistical analysis. Our pipeline provides near seamless integration into existing analysis workflows by allowing substitution and individual execution of each step. The pipeline is available in two versions: an interactive graphical user interface (GUI)-based version for initial exploratory data analysis, and a scalable version using the same software, but streamlined to run with minimal user input for efficient analysis of large datasets. In addition to in-depth single cell-based analysis, basic pixel-based analysis, k-means clustering, visualisation, and pixel-based quantification are provided. All analysis steps can be performed from the container’s command line.

For single-cell based analysis, raw LA-ICP-TOFMS data is processed in HDIP. Isotope channel images are exported as individual 16bit TIFF files for analysis with the MeXpose workflow. Single- or multi-channel TIFF images can be provided from any multiplexed imaging modality. Fiji can be used to pre-process the raw images. The interactive mode allows the user to access the full range of image processing tools in Fiji and to view the effects live on their image set in real-time. The scalable mode provides user-selectable Fiji macros for outlier/hot pixel removal, Gaussian or median filtering for speckle removal, channel stacking, and image tilling. Resulting images are saved as stacks or processed copies of the raw isotope images in 16-bit TIFF format. Cell segmentation requires a 2-channel TIFF stack consisting of a nucleus and a membrane/cytoplasm channel. Cellpose 2.0 is used in both, the interactive and scalable versions of MeXpose to provide cutting-edge cell segmentation performance. Cellpose provides multiple pre-trained deep neural network models based on the Cellpose generalist segmentation algorithm, from which the user can freely select the best performing model^66,67^. In addition, the interactive workflow enables the use of the Cellpose human-in-the-loop pipeline. This feature allows the user to make manual adjustments such as adding/removing object annotations and then retraining the model, resulting in a refined model with improved segmentation performance. By using this approach in conjunction with the Fiji tiling macro, it is possible to make iterative improvements to small subsets of the final data until a satisfactory set of results and a specialist model are obtained. Fine-tuning a model typically requires 3-5 iterations, as recommended in the Cellpose documentation. Successful segmentation results are saved as PNG masks and used to extract morphological features and intensity information from individual cells using Cellprofiler. Similar to pre-processing, the user has access to Cellprofiler functionality in interactive mode. A basic Cellprofiler pipeline file is provided as a template. The user is encouraged to perform the initial setup of a Cellprofiler pipeline either in interactive mode or by running Cellprofiler as standalone software. Marker panels and image channels are subjective to the experiment and require manual configuration. The scalable workflow uses a user pre-configured Cellprofiler pipeline that is executed from the command line. Cellmasks are imported into Cellprofiler along with raw images of all elemental channels of interest. Within Cellprofiler, the user can assign isotopes to image channel names, filter objects that touch the image boundary, filter objects by size, measure object intensities and morphological features, and finally export the data in a tabular format.

MeXpose uses custom Python scripts to perform exploratory data analysis and quantification. These scripts can be deployed either as Jupyter notebooks for interactive analysis or as command line executable scripts for scalable analysis. There are three interactive Jupyter notebooks and three corresponding python scripts provided, a *phenotyping notebook/script* containing all the necessary modules for single-cell data analysis, a *quantification notebook/script* and a *pixel notebook/script* including modules to analyse image data at the pixel level. All scripts can be run independently. The exploratory analysis of the data includes visual inspection using heatmap overlays and histograms, as well as cell phenotyping. Phenotyping is performed by combining unsupervised Phenograph clustering and heatmap-based cluster analysis on scaled and normalised single-cell data. The user can select relevant channels as needed. The heatmap visualisation provides quick identification of relevant cellular phenotypes. A typical analysis workflow would start with the *phenotyping notebook* and later feed cell phenotype-specific data into the *quantification script*. In summary, the structure of the *phenotyping notebook* consists of the following functions:

1. Initial setting and loading of the working directory and single-cell data
2. (Optional) Inspection of raw histograms
3. Size-normalisation (based on ‘real world’ pixel size) and outlier filtering
4. Heatmap visualisation of isotope intensities at the single-cell level (whole sample)
5. Data scaling for accurate clustering
6. Phenograph clustering of the data
7. Dimensionality reduction using UMAP
8. Visualisation of clusters and UMAP embedding
9. Heatmap visualisation of isotope intensities at the single-cell level (per cluster)
10. Data export

The *quantification script* can be run after clustering or as a stand-alone method without clustering, depending on the needs of the user. It uses two input files, a file containing the standard measurements and calibration function, and one or more files containing the single-cell data to be quantified. After setting the working directory and loading all necessary files, the elemental channels for quantification are selected and divided by their corresponding calibration factors. Histograms of the selected channels can then be plotted and saved, and the quantified data can be exported as CSV files.

Pixel-based analysis can be conducted using the *pixel script* by importing a CSV file with intensity information organized in columns, where each row represents a pixel. The user needs to specify the width and height dimensions of the image in pixels and the desired number of clusters for k-means clustering. The resulting image will show each pixel coloured according to its cluster assignment. Using Spearman’s rank correlation coefficients, it is also possible to generate a correlation heatmap for user-selected channels.

### MeXpose image analysis showcased in human skin

Specific to our workflow, we perform the following pre-processing steps for human skin samples: i.) Stacking of nuclei (Ir DNA-Intercalator) and membrane/cytoplasm channels (CD44); ii.) application of a median filter with a pixel size of 1-2 depending on the size of artefacts present; iii.) (optional) splitting image stacks into tiles of ∼300x300 pixels for iterative improvements of the initial segmentation model.

Within Cellpose, the membrane/cytoplasm channel is designated as the primary segmentation channel, with the nuclei channel as an optional secondary channel. The automatic size estimation function is used to determine an appropriate object diameter if no morphological information about the cells is available. To select the initial model, we perform several segmentation runs on a subset of the initial image using the different available models and all default settings. After visual inspection, the most effective model is chosen to perform 3 to 5 iterations of manual annotation and re-training. For this purpose, we use the interactive annotation feature of Cellpose to correct, remove or add annotations to the cells and then retrain the model with our additional annotations. After achieving satisfactory segmentation results on image subsets, we apply the final specialised model to all images of that tissue/cell type.

Cellmasks obtained from Cellpose segmentation are imported into Cellprofiler along with raw images of all relevant channels. In Cellprofiler, potential hot pixels in the raw elemental channel images are first removed using the Smooth Multichannel module^80^ with a *Neighbourhood filter size* of 3.0 and a *Hot pixel threshold* of 50. Next, cell masks are transformed into image objects, and after removing objects that intersect with image boundaries, single-cell elemental intensities and morphological data are exported as a CSV file for downstream analysis.

### Code and data availability

MeXpose code documentation and manuals can be accessed at https://github.com/KoellenspergerLab/MeXpose. The MeXpose docker image is available on Dockerhub at https://hub.docker.com/r/koellenspergerlab/mexpose or can be pulled using ‘*docker pull koellensperger-lab/mexpose:0.1.3’*. All custom python code and scripts are distributed under the MIT licence. The MeXpose docker image is made available under the GPLv3 license. The used data are available upon request. For inquiries contact Gunda Koellensperger at.

## Acknowledgements

The authors acknowledge the financial support from the FG3 Forschungsgruppe (FWF) and from the City of Vienna Fund for Innovative Interdisciplinary Cancer Research (Project no. 21206).

## Author contributions

**G.B.**: Methodology, Software, Writing - Original Draft; **M.S.**: Methodology, Formal Analysis, Investigation, Validation, Visualization, Writing - Original Draft; **P.W.**: Methodology, Investigation; **S.T.**: Writing - Review & Editing; **J.Z.**: Supervision, Writing - Review & Editing; **L.W.**: Investigation, Writing - Review & Editing; **N.F.**: Supervision, Funding Acquisition; Writing - Review & Editing; **G.K.**: Supervision, Funding Acquisition, Writing - Original Draft

## Ethics declarations

The authors declare no competing interests.

