## Supplementary Information for "MeXpose - A modular imaging pipeline for the quantitative assessment of cellular metal bioaccumulation"

* Corresponding author:

Gunda Koellensperger

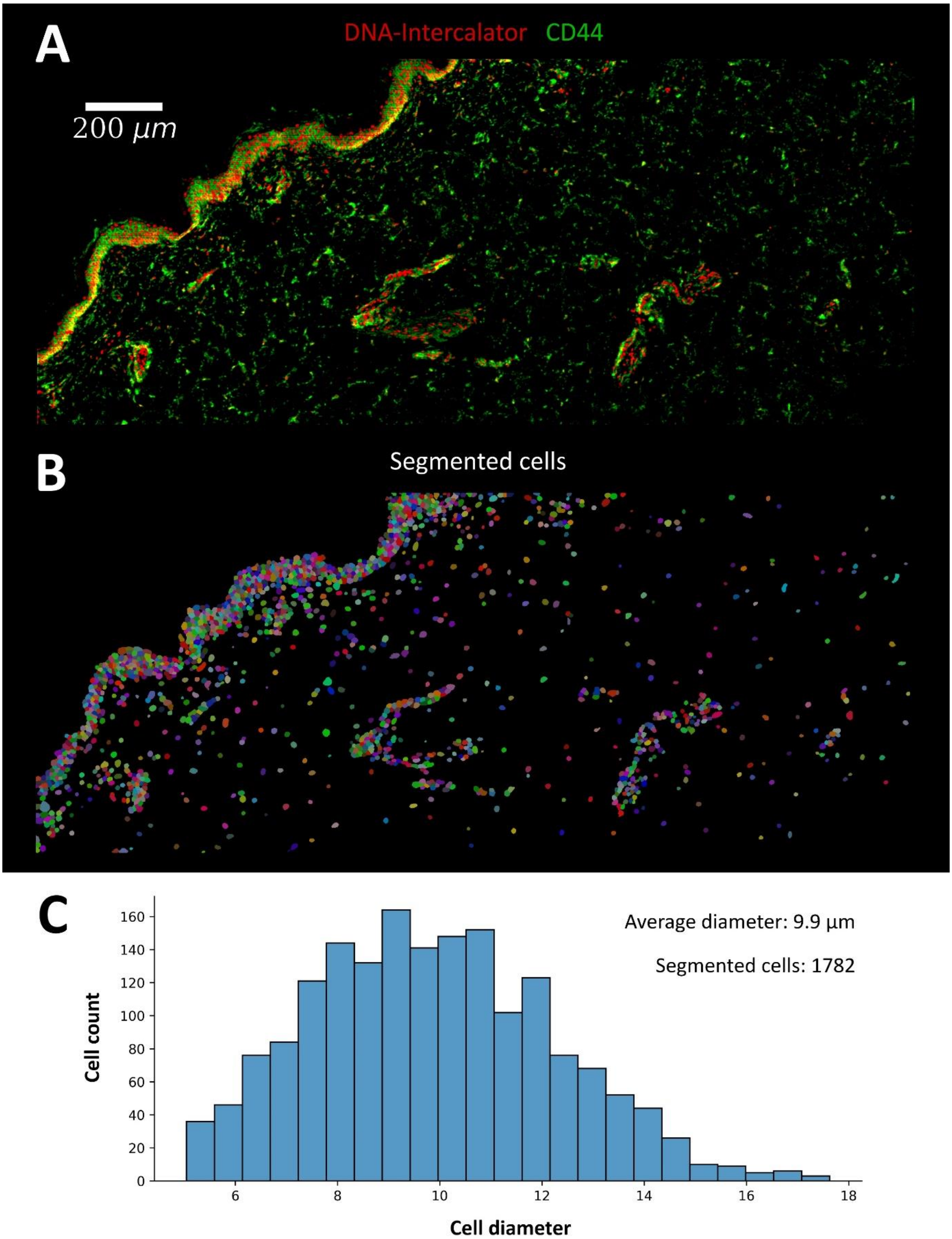


**Supplementary Figure 1**: (A) Overlay images of cell membrane (CD44) and nuclei marker (DNA-Intercalator) in human skin as measured by LA-ICP-TOF-MS (1µm spatial resolution, 200 Hz ablation speed). These channels were used for segmentation following stacking and pre-processing. (B) Map of cellular objects (multidimensional data with characterized area and intensity)- (C) Size distribution of the cellular objects.  A total of 1782 cells were segmented in the image with an average diameter of 10 µm.

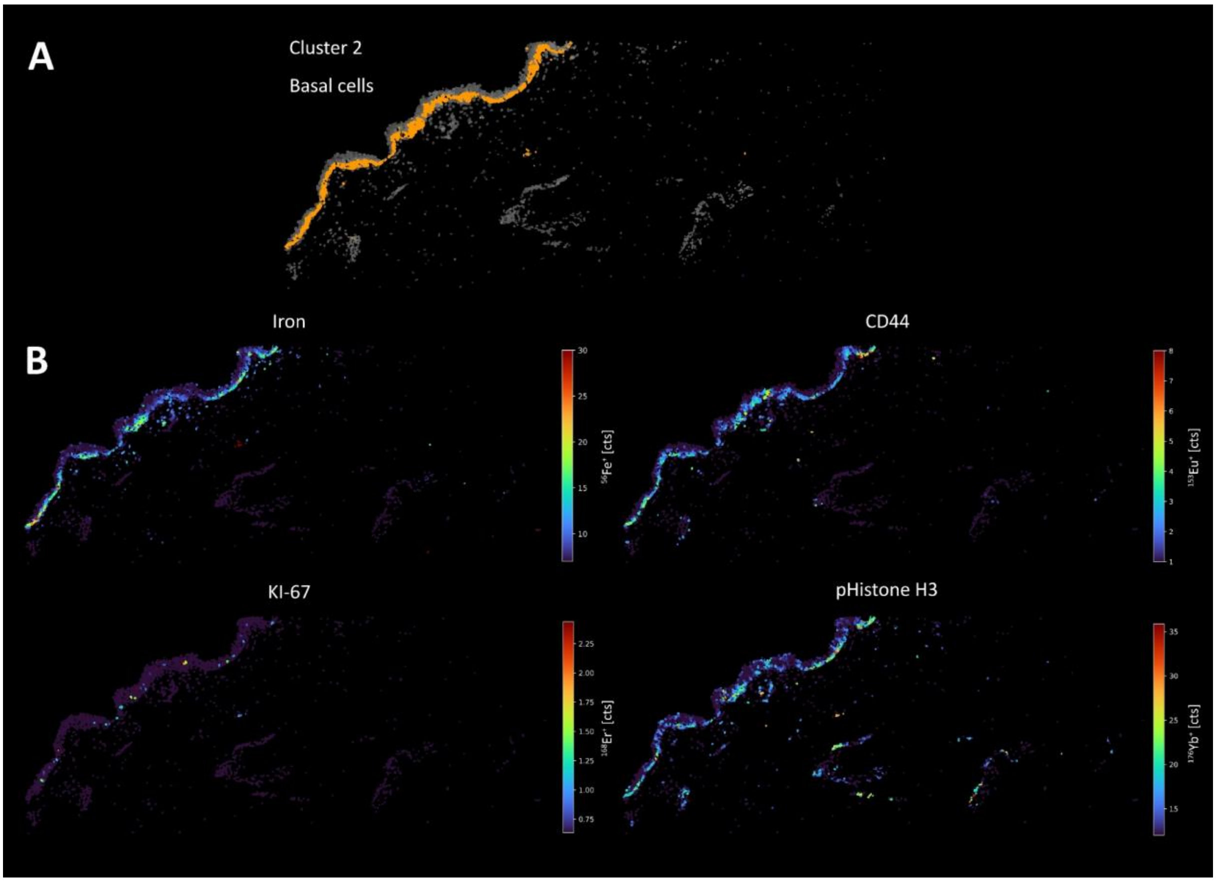


**Supplementary Figure 2**: (A) Visualisation of cluster 2 (basal cells) within all of the segmented cells. (B) Iron showed a distribution characteristic of basal cells and was therefore helpful in phenotyping cluster 2, in addition to CD44, KI-67 and pHistone H3.

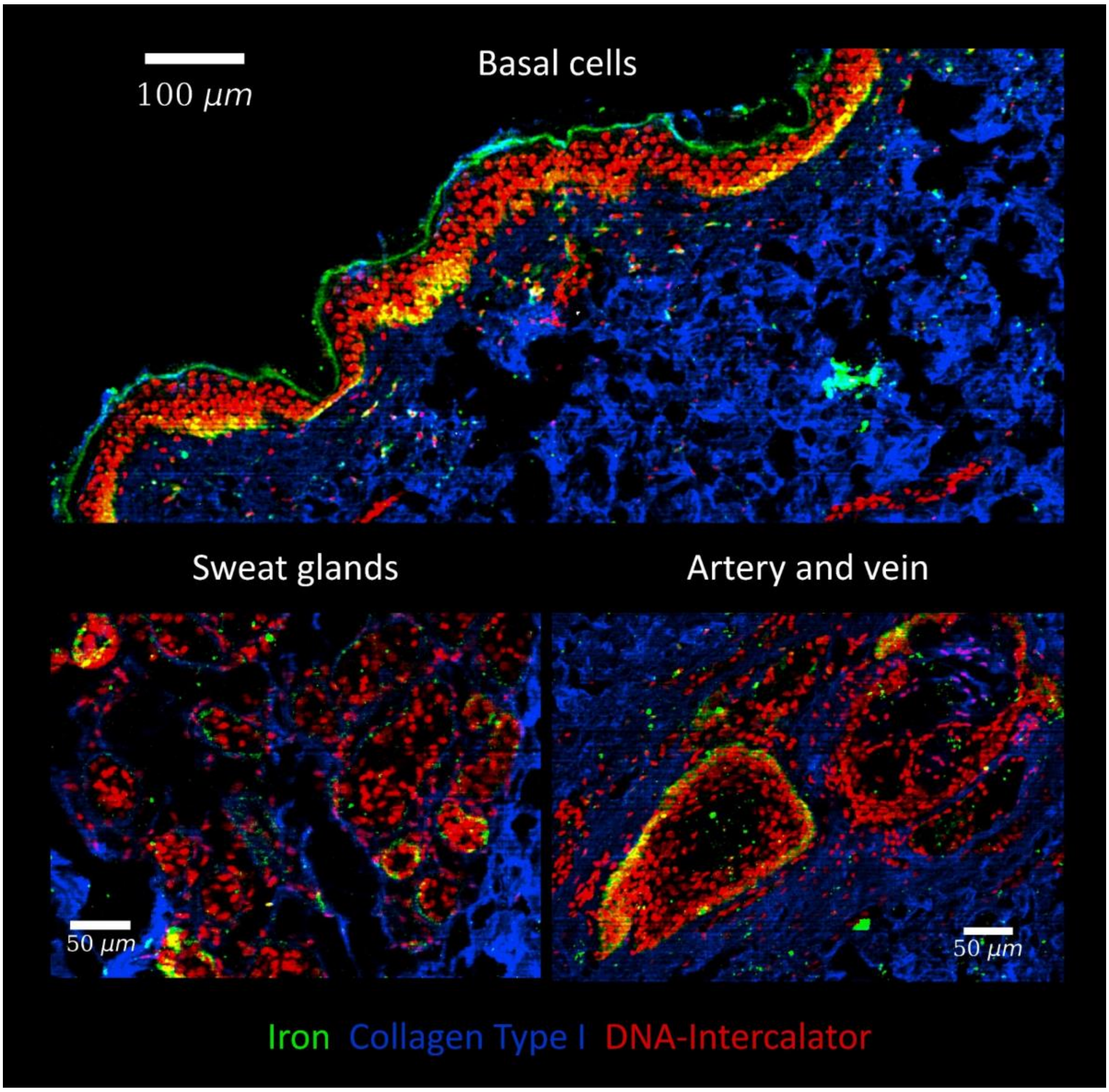


**Supplementary Figure 3**: Signal overlay of iron (green), collagen type I (blue) and DNA-intercalator (red) from different skin regions. Iron shows characteristic distributions in structural features of the skin such as basal cells, sweat glands or arteries even after FFPE treatment of the skin, which can improve phenotype clustering.

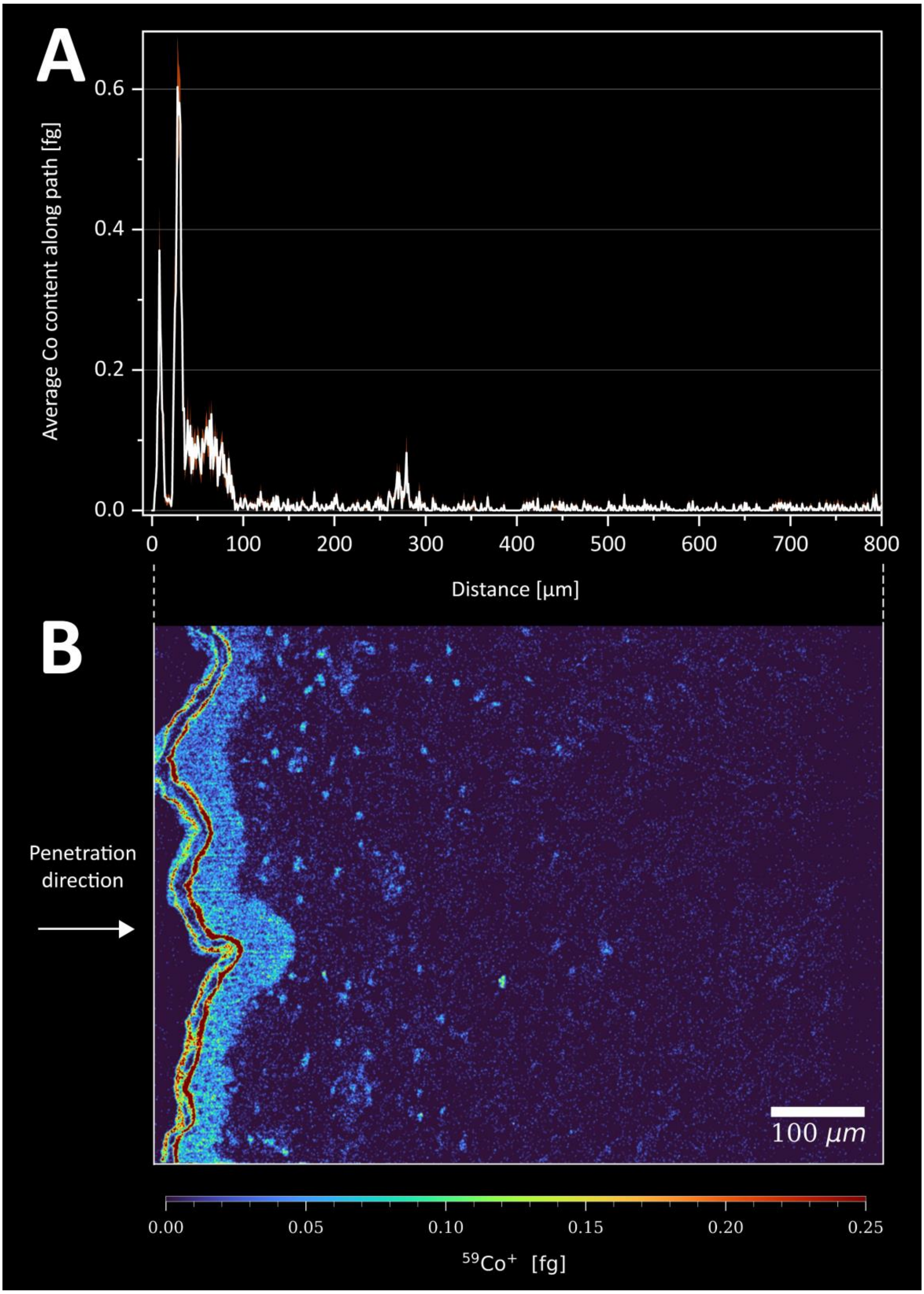


**Supplementary Figure 4:** (A) Average amount of cobalt along a path plotted against the distance in the tissue. (B) shows the section on which the plot is based (LA-ICP-TOFMS analysis, 1 µm pixel size, 200 Hz pixel acquisition rate). The whole area has been integrated from left to right. The permeation mainly occurs 100 µm into the skin, i.e. into the epidermis. Only a small fraction of the cobalt is able to penetrate further into the dermis.

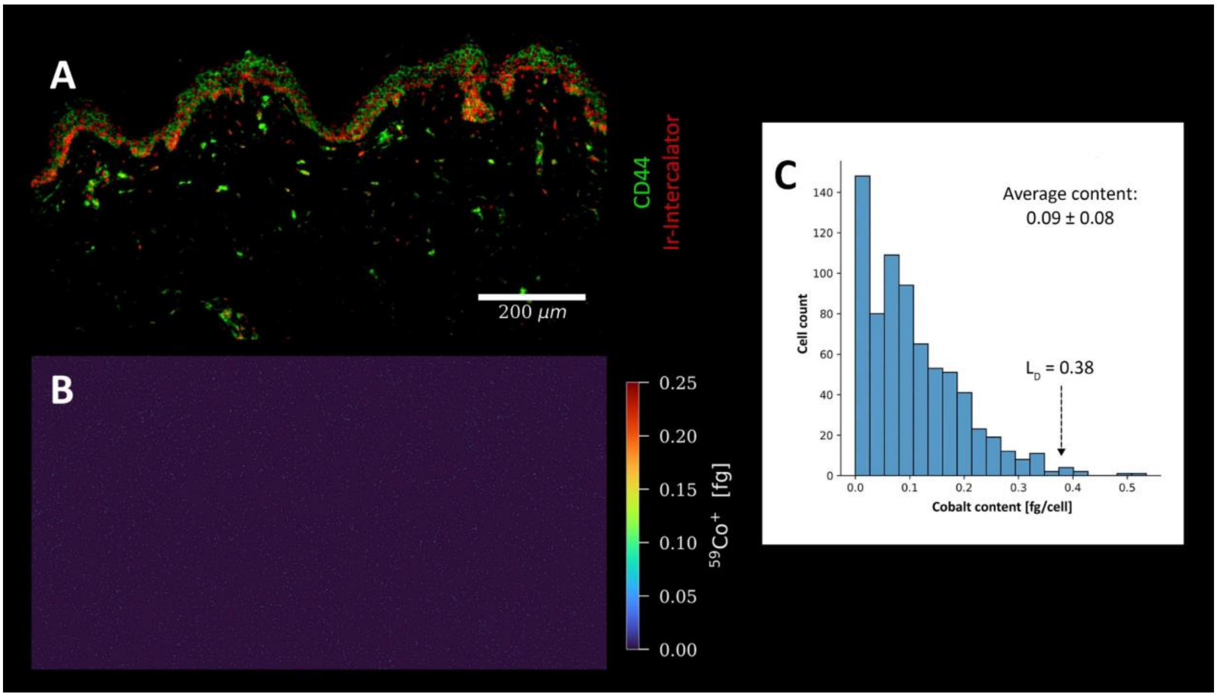


**Supplementary Figure 5**: (A) Signal overlay of CD44 (green) and DNA intercalator (red) to visualise cells in a control sample (tagged with metal-conjugated antibodies, incubation with H2O). The cobalt distribution in (B) shows that there is no measurable amount of cobalt in the tissue. There is only a background noise spread over the whole image. (C) shows the histogram for segmented cells with an average content of 0.1 fg. The LOD obtained is higher than 99% of the cells present in the control sample.

#### Quantitative analysis calls for validation (Supplementary Figure 6)

Whether the applied histochemistry allows for simultaneous phenotyping and cellular metal enrichment needs to be evaluated for each specific exposure study. As in any quantitative study, additional experiments are required proving the method fit for purpose. The impact of the sample preparation is investigated on consecutive thin sections using pixel-based data, as cell-based data in tissue are not amenable without staining procedures. Therefore, the quantitative module of MeXpose, includes a script for pixel-based image analysis. Supplementary Figure 5 shows the Co distribution in native skin of consecutive sections. K-means clustering for three expected clusters (background, the cornified layer and cells) showed for all three clusters an excellent agreement– both in terms of Co quantification and location- when comparing consecutive sections with and without staining.

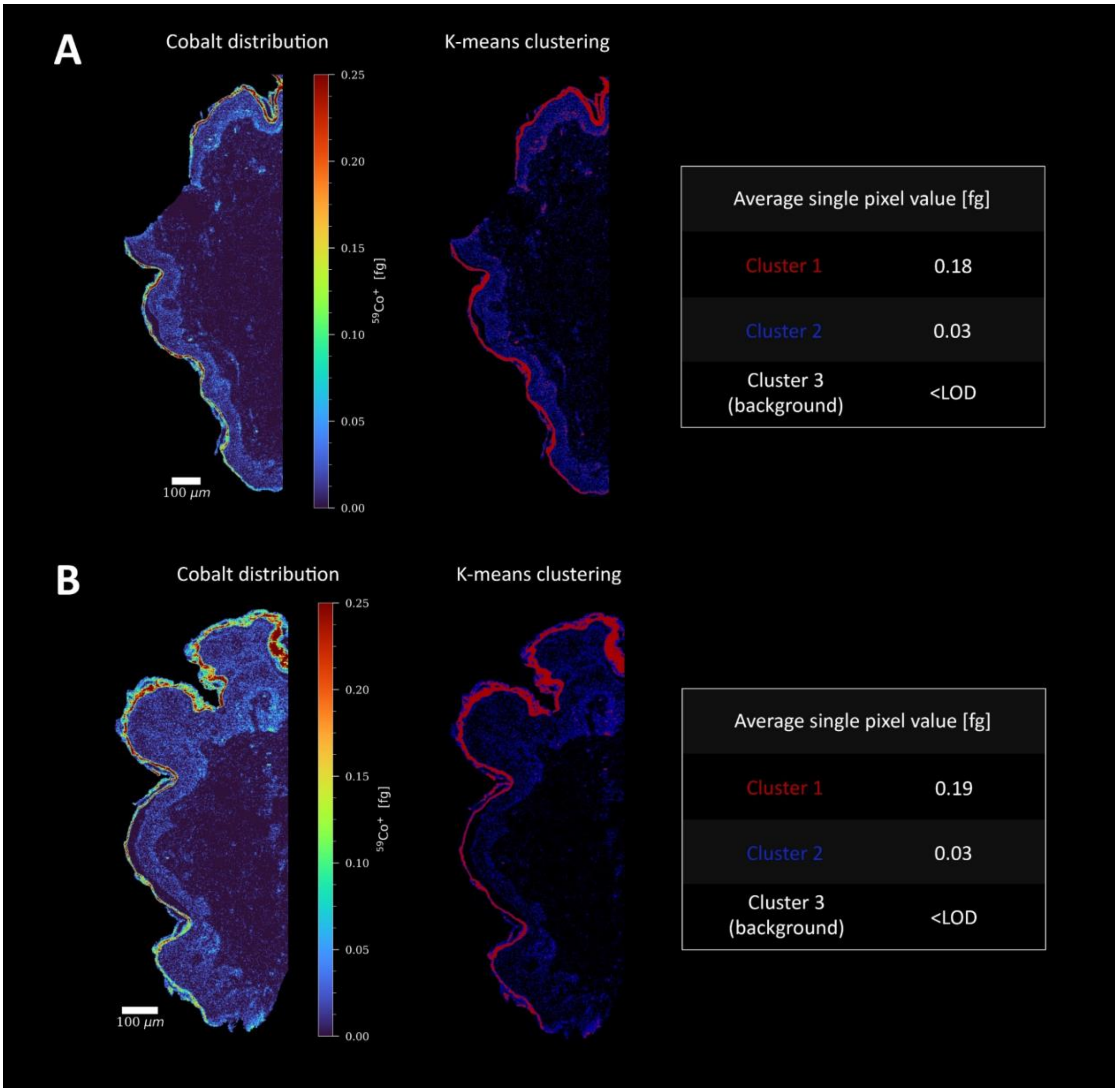


**Supplementary Figure 6**: Co distribution in consecutive sections, (A) unlabelled and (B) labelled, showing the calculated Co amounts obtained by k-means clustering for the three user-defined clusters (background, cornified layer, cells).

### Null hypothesis using Mann-Whitney U test (Supplementary Figure 7)

The single cell Co accumulation, obtained for each cell cluster, respectively (Figure 4E), was tested for significant differences by statistical analysis. By using the Mann-Whitney U test a p-value matrix was generated allowing pair-wise comparison (Fig. S7). Cluster 0, characterised by the highest cell number, showed a notably low cobalt concentration with an average of 1.2 fg per cell. There were no significant differences between clusters 3, 4 and 6, with mean contents of 2.1-2.2 fg per cell. Similarly, with an average of 2.8 fg per cell, clusters 1 and 5 showed considerable similarity. Notably, cluster 2 showed significant differences from all clusters (including a remarkable p-value of 4.22E-91 for cluster 0) and stood out with 3.8 fg per cell.

| | **Cluster 0** | **Cluster 1** | **Cluster 2** | **Cluster 3** | **Cluster 4** | **Cluster 5** | **Cluster 6** |
|---|---|---|---|---|---|---|---|
| **Cluster 0** | 1 | 4.07E-74 | 4.22E-91 | 4.29E-19 | 6.96E-22 | 2.05E-21 | 1.53E-11 |
| **Cluster 1** | 4.07E-74 | 1 | 7.46E-15 | 9.05E-16 | 2.78E-10 | 1.66E-01 | 1.93E-09 |
| **Cluster 2** | 4.22E-91 | 7.46E-15 | 1 | 7.38E-33 | 8.96E-25 | 1.47E-07 | 4.97E-18 |
| **Cluster 3** | 4.29E-19 | 9.05E-16 | 7.38E-33 | 1 | 1.35E-01 | 2.40E-04 | 8.46E-01 |
| **Cluster 4** | 6.96E-22 | 2.78E-10 | 8.96E-25 | 1.35E-01 | 1 | 7.23E-03 | 2.47E-01 |
| **Cluster 5** | 2.05E-21 | 1.66E-01 | 1.47E-07 | 2.40E-04 | 7.23E-03 | 1 | 2.14E-03 |
| **Cluster 6** | 1.53E-11 | 1.93E-09 | 4.97E-18 | 8.46E-01 | 2.47E-01 | 2.14E-03 | 1 |

**Supplementary Figure 7**: Significant differences between clusters were assessed using p-values determined by the Mann-Whitney U-test. Using α = 0.05 as the significance level, significant values (<0.05) were highlighted green and non-significant values (>0.05) red.

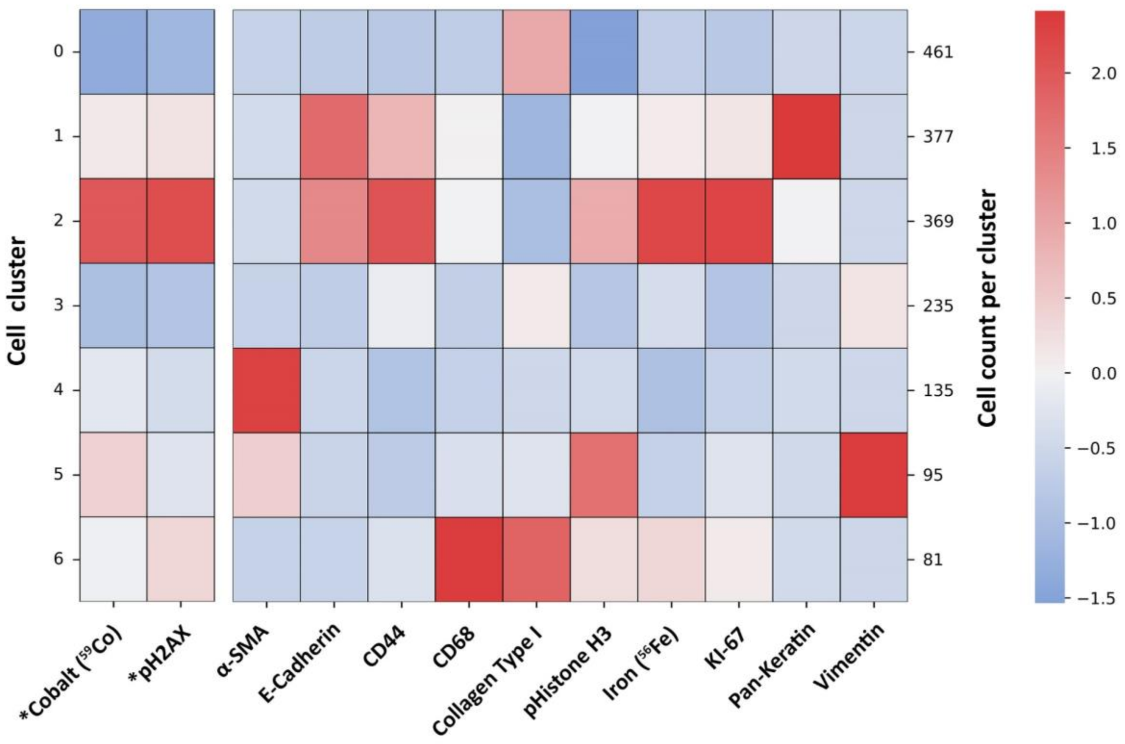


**Supplementary Figure 8**: Heat map plotting cobalt and pH2AX along the heat map of Figure 3. As these read-outs are not characteristic of the phenotypes, they have not been included in the clustering itself. The high intensity of cobalt and DNA damage in cluster 2 can be clearly seen.

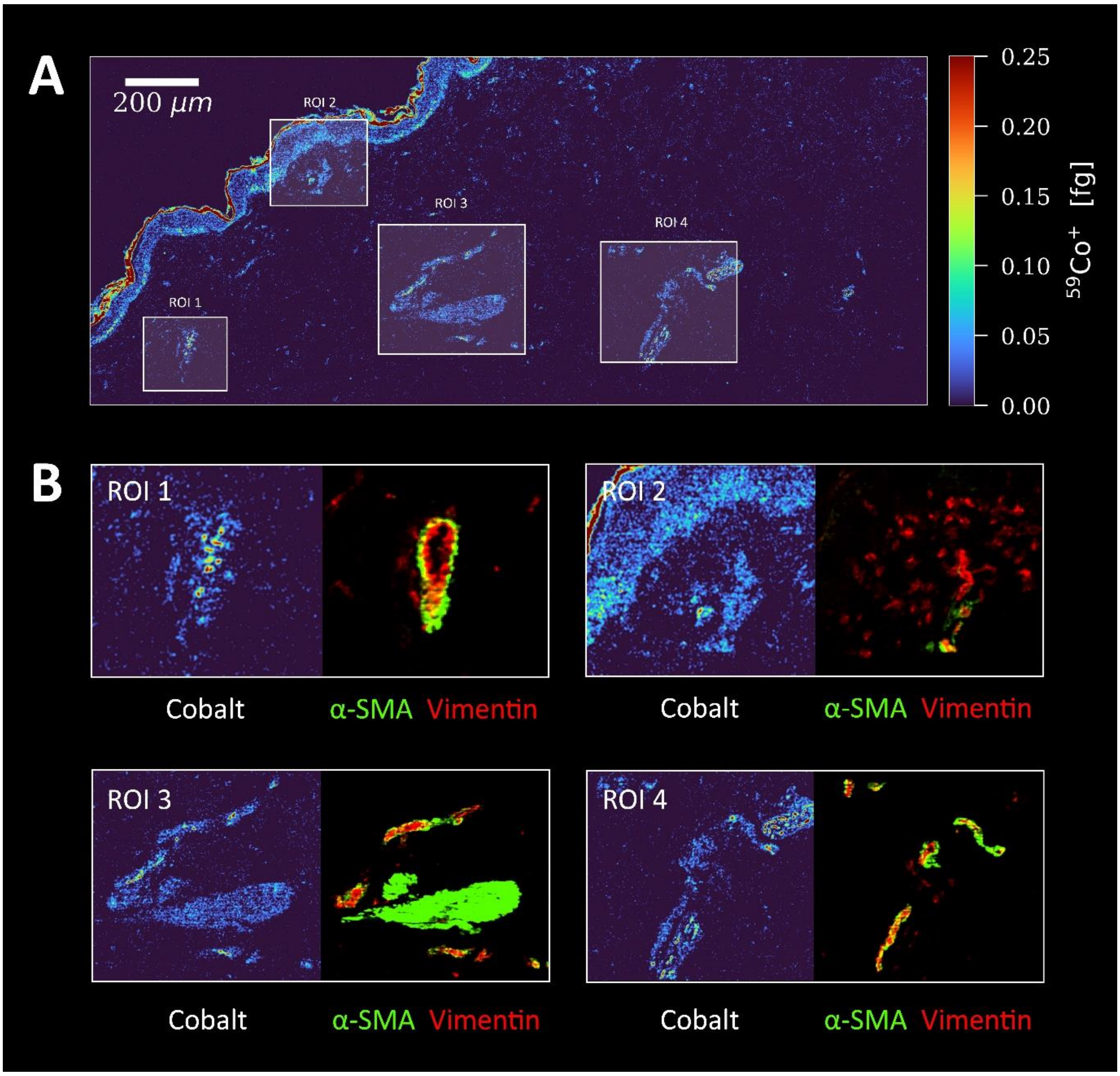


**Supplementary Figure 9:** (A) Quantitative imaging of cobalt in human skin (LA-ICP-TOFMS analysis, 1 µm pixel size, 200 Hz pixel acquisition rate). Regions of interest (ROI) for blood vessels are outlined in the tissue section. (B) Cobalt distribution compared with the location of blood vessels visualised by a-SMA (green) and vimentin (red). Increased accumulation in blood vessels is indicated.

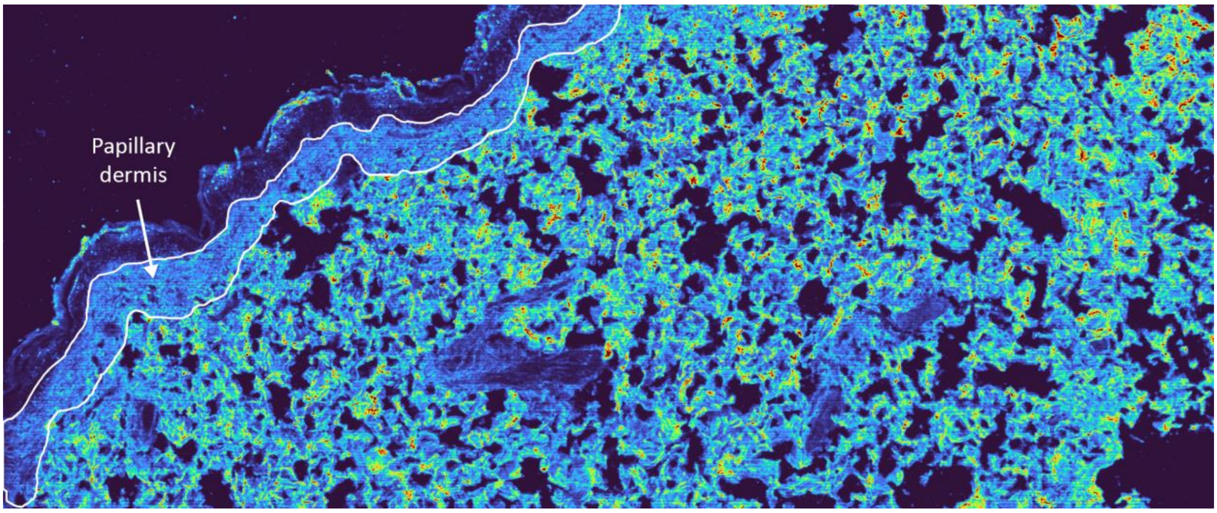


**Supplementary Figure 10:** Signal intensity map of collagen type I ($^{169}$Tm) for human skin (LA-ICP-TOFMS analysis, 1 µm pixel size, 200 Hz pixel acquisition rate). Papillary dermis (outlined in white) can be distinguished from reticular dermis by collagen expression. While collagen fibers in the papillary dermis are thin and loosely arranged, those in the reticular dermis are thicker and densely packed.

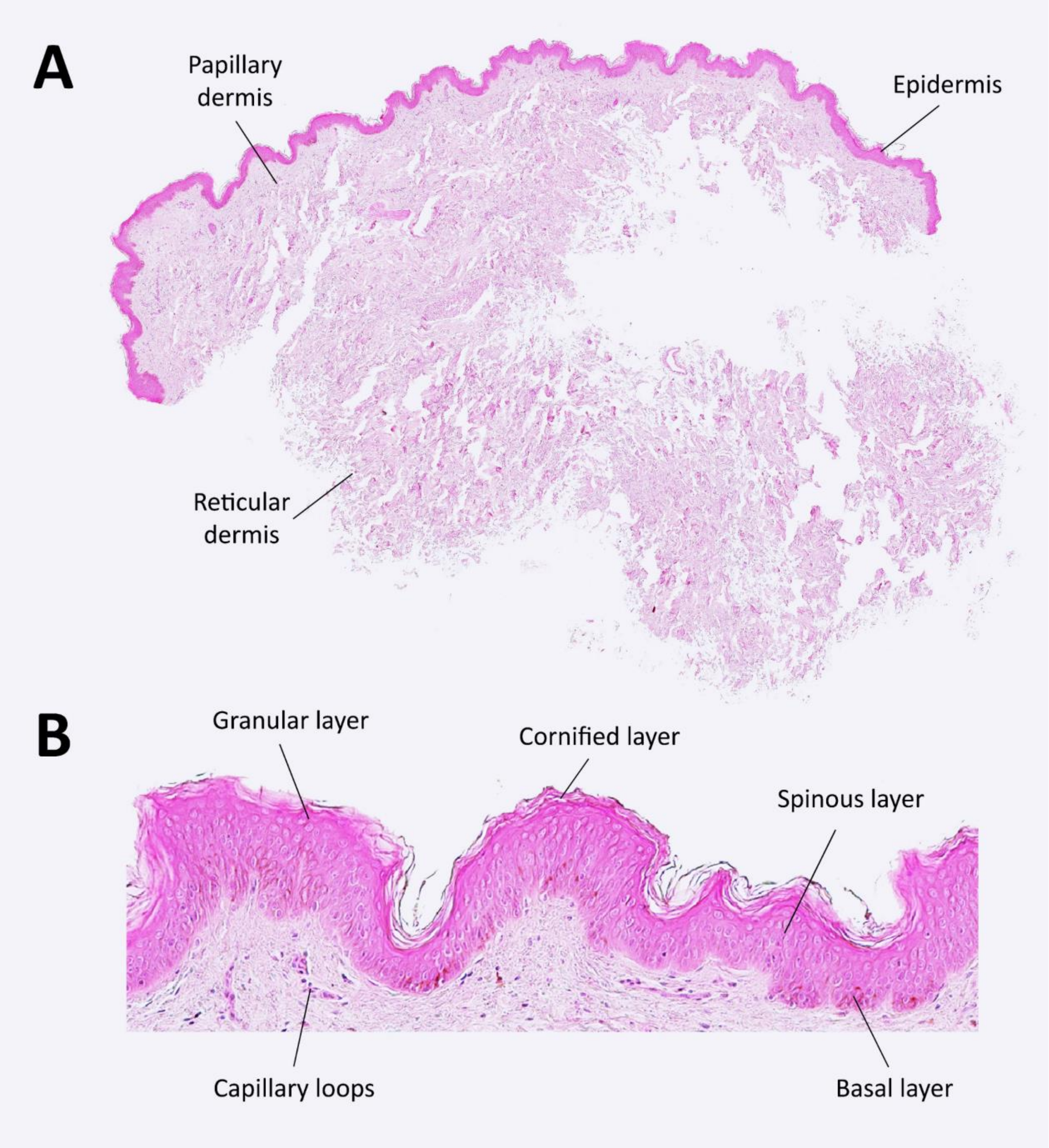


**Supplementary Figure 11:** (A) Tissue section of NativeSkin stained with hematoxylin and eosin (H&E). The epidermis and papillary dermis were consistently observed in their entirety, although there were instances of incomplete reticular dermis due to the sectioning process. (B) provides a close-up of the epidermis for detailed examination.

**Supplementary Table 1**: List of metal-conjugated antibodies

| Antibody target | Clone | Metal tag | Catalog number | Target |
|---|---|---|---|---|
| **α-SMA** | 1A4 | $^{141}Pr$ | 3141017D | Smooth muscle cells |
| **Vimentin** | D21H3 | $^{143}Nd$ | 3143027D | Mesenchymal cells, fibroblasts |
| **Pan-Keratin** | C11 | $^{148}Nd$ | 3148020D | Cornified layer/ keratinocytes |
| **CD44** | IM7 | $^{153}Eu$ | 3153029D | Hyaluronic acid receptor /intercellular adhesion |
| **E-Cadherin** | 24E10 | $^{158}Gd$ | 3158029D | Epithelial cell membrane |
| **Collagen Type I** | Polyclonal | $^{169}Tm$ | 3169023D | Dermal collagen fibers |
| **CD68** | KP1 | $^{159}Tb$ | 3159035D | Macrophages |
| **pH2AX** | S139 | $^{165}Ho$ | 3165036D | DNA damage |
| **KI-67** | B56 | $^{168}Er$ | 3168022D | Proliferation |
| **pHistone H3** | HTA28 | $^{176}Yb$ | 3176024D | Mitosis |

**Supplementary Table 2**: LA-ICP-TOFMS parameters

| **ICP-TOFMS** | |
|---|---|
| RF Power [W] | 1440 |
| Sampling depth [mm] | 3.5 |
| Cone materials | Ni |
| Plasma gas flow [L $min^{-1}$] | 14 |
| Auxiliary gas flow [L $min^{-1}$] | 0.80 |
| Nebulizer gas flow [L $min^{-1}$] | 1.06 |
| Measurement mode | Collision cell technology (CCT) |
| CCT gas | 93% He *(v/v)*, 7% $H_2$ *(v/v)* |
| CCT gas flow [mL $min^{-1}$] | 4.2 |
| m/z range | 14-256 |
| **Laser ablation** | |
| Spot size | 2 µm (circular) |
| Interspacing | 1 µm |
| Repetition rate | 200 Hz |
| Dosage | 2 |
| Fluence | 0.4 - 1.0 J $cm^{-2}$ |
